# Characterization of the utility of three nebulizers in investigating infectivity of airborne viruses

**DOI:** 10.1101/2021.03.11.435057

**Authors:** Sadegh Niazi, Lisa K. Philp, Kirsten Spann, Graham R. Johnson

**Affiliations:** Queensland University of Technology (QUT), Faculty of Science, School of Earth and Atmospheric Sciences, Brisbane 4001, Australia; Queensland University of Technology (QUT), Faculty of Health, School of Biomedical Sciences, Brisbane 4001, Australia; Queensland University of Technology (QUT), Faculty of Health, School of Biomedical Sciences, Centre for Immunology and Infection Control, Brisbane 4001, Australia

## Abstract

Laboratory-generated bioaerosols are widely used in aerobiology studies of viruses, however few comparisons of alternative nebulizers exist. We compared aerosol production and virus survival for a Collison nebulizer, vibrating mesh nebulizer (VMN), and hydraulic spray atomizer (HAS). We also measured the dry size distribution of the aerosols produced, calculated the droplet sizes before evaporation and the dry size distribution from normal saline solution. Dry count median diameters of 0.25, 0.63 and 0.76 µm were found for normal saline from the Collison nebulizer, VMN and HSA, respectively. The volume median diameters were 2.91, 3.2 and 2.43 µm, respectively. The effect of nebulization on the viability of two influenza A viruses (IAVs) (H1N1, H3N2) and human rhinovirus (HRV)-16, was assessed by direct nebulization into an SKC Biosampler. The HSA had least impact on surviving fractions (SFs) of H1N1 and H3N2 (89±5%, 94±3%), followed by the Collison nebulizer (82±2%, 82±3%). The VMN yielded SFs of 78±2% and 76±2%, respectively. Conversely, for HRV-16, the VMN produced higher SFs (86±15%). Our findings indicate that although the VMN had the greatest impact on IAV survival, it produced higher aerosol concentrations within the airborne-size range making it more suitable where high aerosol mass production is required.

**Importance:** Viral respiratory tract infections cause millions of lost days of work and physician visits globally, accounting for significant morbidity and mortality. Respiratory droplet and droplet nuclei from infected hosts are the substantial potential carriers of such viruses within indoor environments. Laboratory-generated bioaerosols are applied in understanding the transmission and infection of viruses, simulating the physiological aspects of bioaerosol generation in a controlled environment. However, little comparative characterization exists for nebulizers used in infectious disease aerobiology, including Collison nebulizer, Vibrating mesh nebulizer, and hydraulic spray atomizer. This study characterized the physical features of aerosols generated by laboratory nebulizers, and their performance in producing aerosols at a size relevant to airborne transmission used in infectious disease aerobiology. We also determined the impact of nebulization mechanisms of these nebulizers on the viability of human respiratory viruses, including IAV H1N1, IAV H3N2 and HRV-16.

## Introduction

Respiratory viruses are responsible for significant morbidity and mortality as well as millions of lost days of work and physician visits globally (1, 2). Evidence suggests that the airborne mode of transmission plays a significant role in the spread of respiratory viruses (3, 4). Respiratory droplets and droplet nuclei generated from the human host respiratory system are potential carriers of pathogens within indoor environments (5, 6). However, there is no consistent explanation that fully addresses the ability of these viruses to survive in an airborne state (7). It has been reported that a combination of environmental and biological factors can affect the efficiency of airborne transmission of viruses in indoor environments (8). In modelling airborne transmission, laboratory-generated pathogen-laden aerosols are broadly used to investigate the transmission, infection and toxicology of respiratory viruses (8-10). Techniques used to generate virus-laden aerosols in the laboratory enable greater control over aerosol characteristics, including their droplet concentration, size and efficacy for carrying viruses (11). Animal models have also been used to investigate the infection course and pathogenesis of inhaled microorganisms (12). Here, the route of exposure, aerosol size, and infectious dose can influence the infection development and pathogenic outcomes. In these models, the method of aerosolization is critical, as reduced virus viability due to mechanical preparation would influence study outcomes. Considering this critical factor, the field lacks a comprehensive study that fully characterizes the physical and biological properties of carrier aerosols generated by laboratory-use nebulizers, particularly for the simulation of respiratory virus-laden aerosols in a clinical environment.

The Collison nebulizer, as the gold standard, has dominated bioaerosol generation research since its invention in 1932 (13-15). It applies Bernoulli’s theory, impaction of a liquid suspension against the interior of a glass to generate small size aerosols (16). The type of liquid suspension, its viscosity and surface tension, are the principal factors that influence the droplet size and concentration of aerosols generated by a Collison nebulizer (17). Within the literature reporting Collison nebulizer generated aerosols, studies utilize different nebulization solution viscosities, resulting in the multiple size distributions reported from Collison use. It has also been hypothesized that impaction, shear forces and recirculation of an infectious sample within the Collison nebulizer may damage microorganisms, potentially decreasing pathogen viability or infectivity (18, 19). However, there is no available well-designed studies to test these effects on pathogen viability.

The vibrating mesh nebulizer (VMN), has been introduced more recently as a high performance treatment delivery system for patients with respiratory diseases (20, 21). VMNs employ electroformed plates that vibrate to generate aerosols (22). Two types of VMN, passive and active, have been commercially deployed. An example of a passive VMN is the Omron MicroAir NE-U22, which contains a perforated vibrating plate underlying the fluid reservoir with around 6000 tapered holes of 3 µm diameter. Alternatively, the Aeroneb Pro nebulizer is an active VMN, utilizing a micropump system which delivers fluid to a vibrating plate containing up to 1000 dome-shaped apertures. In both systems, aerosols are generated by applying an alternating electric potential to a piezoelectic, triggering the mesh to move back and forth by a few micrometres to aerosolize liquids brought into contact with the mesh surface (17). An alternative to the Collison and VMN nebulization, is the hydraulic spray atomizer (HSA). The HSA concentrates the suspension into a stream by forcing it through a very small hole. There is a one-way valve in the nozzle that maintains air from flowing back into the pump and allows for suction within the pump so that liquid can be pulled up the tube. However, these latter two types of nebulizer VMN and HSA have rarely been applied in the infectious disease aerobiology research field. In this study, we hypothesized that a VMN or an HSA is superior to the commonly used Collison nebulizer, as they result in less mechanical stress on respiratory viruses and therefore enable a higher viable dose to be delivered for experimental purposes.

This study was therefore designed to determine the influence of nebulization methods of the 1-jet Collison nebulizer, VMN and HSA on the viability of human respiratory viruses Influenza A virus (IAV) H1N1, IAV H3N2 and Human rhinovirus-16 (HRV-16). We also characterized the size and weight of aerosols produced by these three nebulizers.

## Results

### Aerosol Output and Size Analysis

**Figure 1**. shows the lognormal fit to ASD normalised with respect to the maximum sizing channel concentration of the dried aerosols generated by the Collison nebulizer, VMN and HSA loaded with 0.05 g L^-1^ NaCl solution as measured with SMPS. Initial droplet size (dotted curves) of these aerosols before evaporation were calculated using ***Equation 1*** and finally the ASDs were calculated for a solution composed of 9 g L^-1^ NaCl (dashed curves). **Table 1** summarizes the detailed properties of the aerosols produced. The Collison generated smaller aerosols compared to the other two nebulizers, with count median diameters (CMD) of 0.045 µm and 0.25 µm for 0.05 g L^-1^ and 9 g L^-1^ NaCl solutions, respectively. However, the corresponding volume median diameters (VMD) of these two NaCl solutions were 0.50 µm and 2.91 µm. VMN generated larger aerosols than the Collison with a CMD of 0.11 µm and 0.63 µm for 0.05 g L^-1^ and 9 g L^-1^ NaCl solutions, and VMD of 0.56 µm and 3.2 µm, respectively. HSA generated the largest aerosols with CMD of 0.13 µm and 0.76 µm, although the VMD were 0.43 µm and 2.43 µm, for 0.05 g L^-1^ and 9 g L^-1^ NaCl solutions, respectively. The geometric standard deviations (GSDs) of the aerosols produced by the Collison, VMN and the HSA were 2.47, 2.09 and 1.86, respectively, indicating that HSA generated a more uniform aerosol size compared to the other two nebulizers.

**Figure 1.**
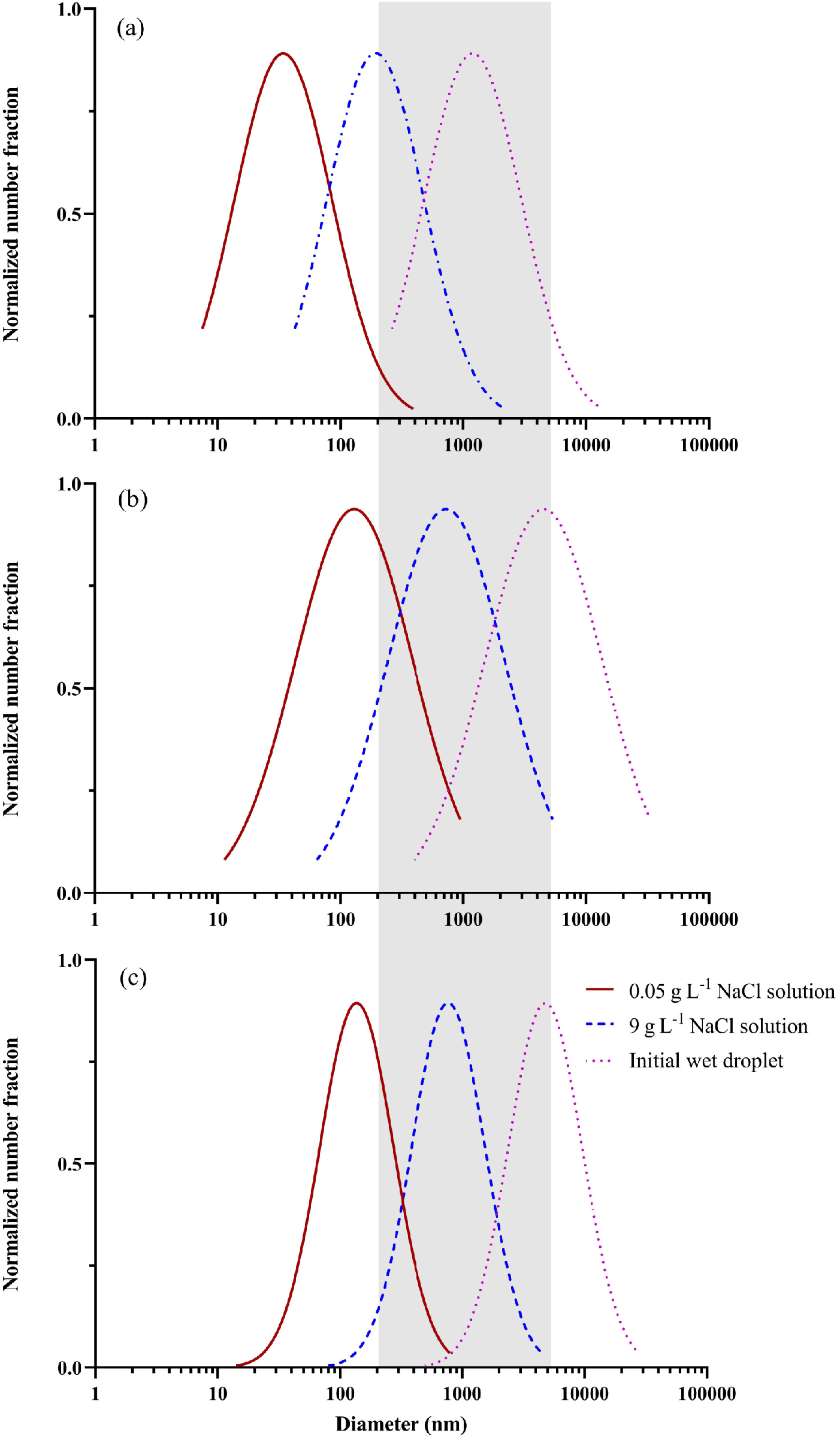
Dried aerosols size distributions for aerosols generated by (a) Collision, (b) VMN and (c) HSA loaded with 0.05 g L^-1^ NaCl (solid lines), their initial droplet sizes (dotted curves) and for a solution of 9 g L^-1^ NaCl (dashed curves). The shading indicates the usable size (the size can carry respiratory viruses and is in the airborne size range), suitable for studying infectious diseases in the aerobiology research field.

**Table 1.**
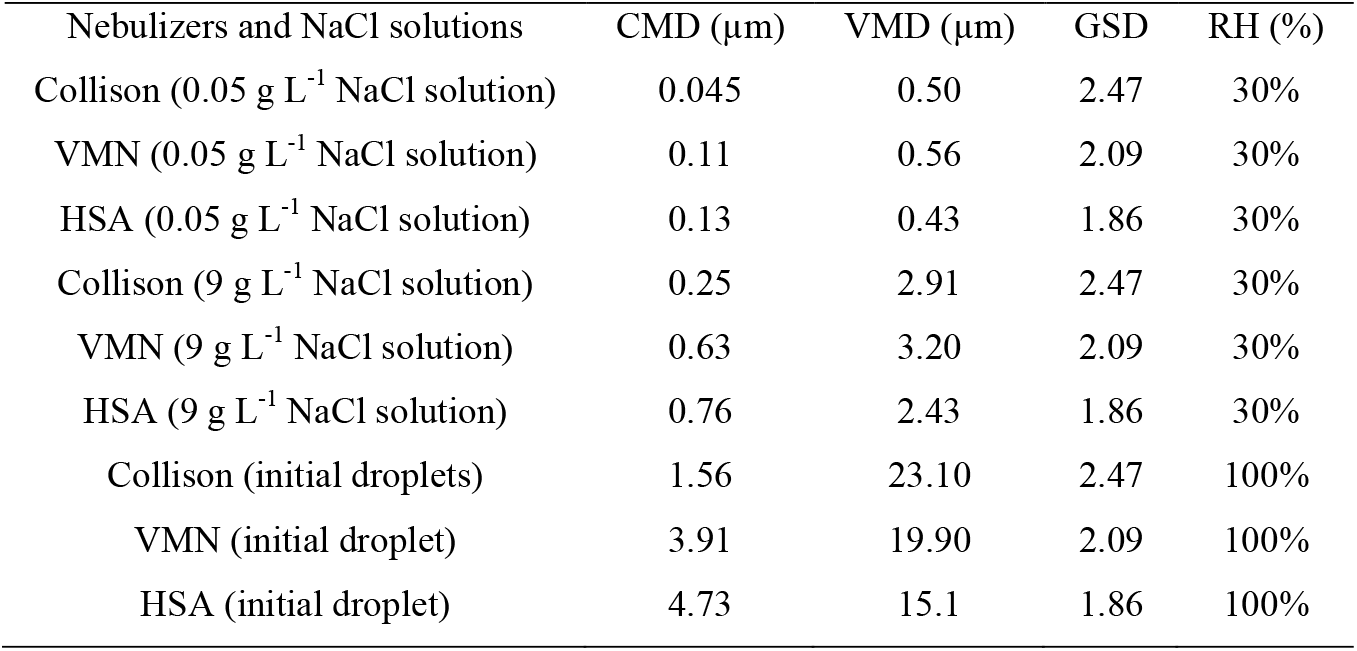
The properties of dry and wet aerosols generated by Collison nebulizer, VMN and HAS derived from two NaCl solutions.

### Viral Surviving Fractions Post-Aerosolization

Comparison of SFs for IAVs H3N2 and H1N1 and HRV-16 for Collison, VMN and HSA were performed by nebulization of viral suspension directly into an operating SKC cup and results are presented in **Figure 2**. The average SFs of IAV H1N1 for the Collison, VMN and HSA were 0.82±0.02, 0.78±0.02 and 0.89±0.05, respectively, while the average SFs for IAV H3N2 were 0.82±0.03, 0.76±0.02 and 0.94±0.03. The corresponding average SFs of HRV-16 after running the Collison, VMN and HSA were 0.83±0.08, 0.86±0.15 and 0.85±0.06, showing that these nebulizers had little effect on HRV-16 survival compared to that of survival of IAV strains. Although HSA is slightly better for IAVs, One-way ANOVA demonstrated no statistic differences in SFs of the tested viruses between three mentioned nebulizers.

**Figure 2.**
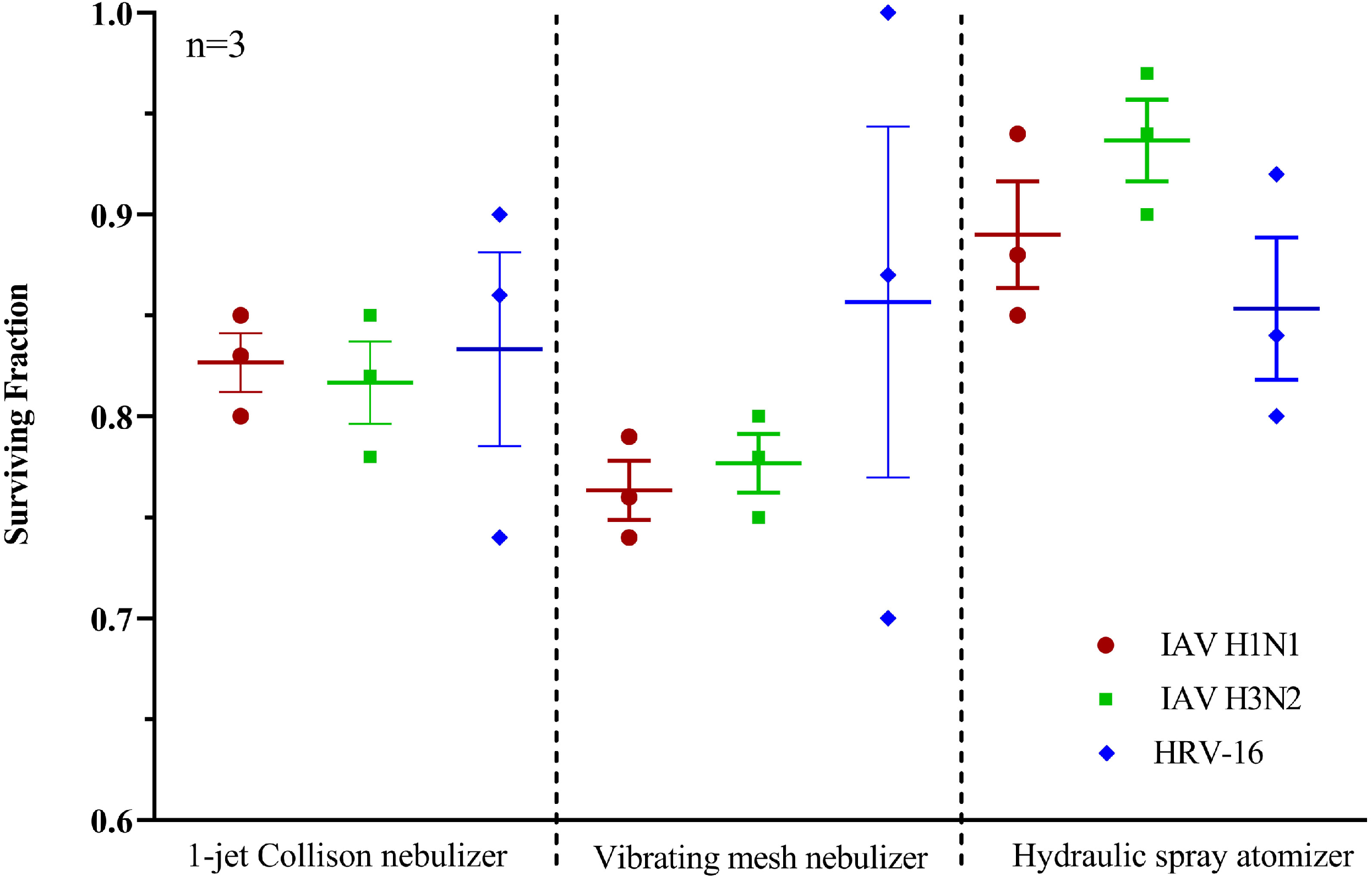
Surviving fraction (calculated based on *equation 2*) of IAV H1N1, IAV H3N2 and HRV-16 immediately post-nebulization. Experiments were conducted in triplicate.

We also tested the shear and impact forces delivered by the high-velocity air streams of the 1-jet Collison nebulizer on the viability of viruses suspended during 30 min run times **(Figure 3)**. There was a gradual decrease in SFs for all three viruses over 30 min. SFs of IAVs H1N1 and H3N2 declined from 0.82±0.02 at 5 min to 0.67±0.03 at 30 min, and 0.82±0.03 at 5 min to 0.68±0.02 at 30 min, respectively, these differences between 5 and 30 min operating times were statistically significant (*P*=0.003 and *P*=0.012, respectively), suggesting that the Collison may not be an appropriate option for studies that require longer nebulizer operating time or high concentration of viable virus in aerosols. However, the SF of HRV-16 decreased less, from 0.83±0.08 at 5 min to 0.73±0.05 at 30 min, which was not significantly different, suggesting that HRV is less susceptibility to mechanical stress compared to IAVs.

**Figure 3.**
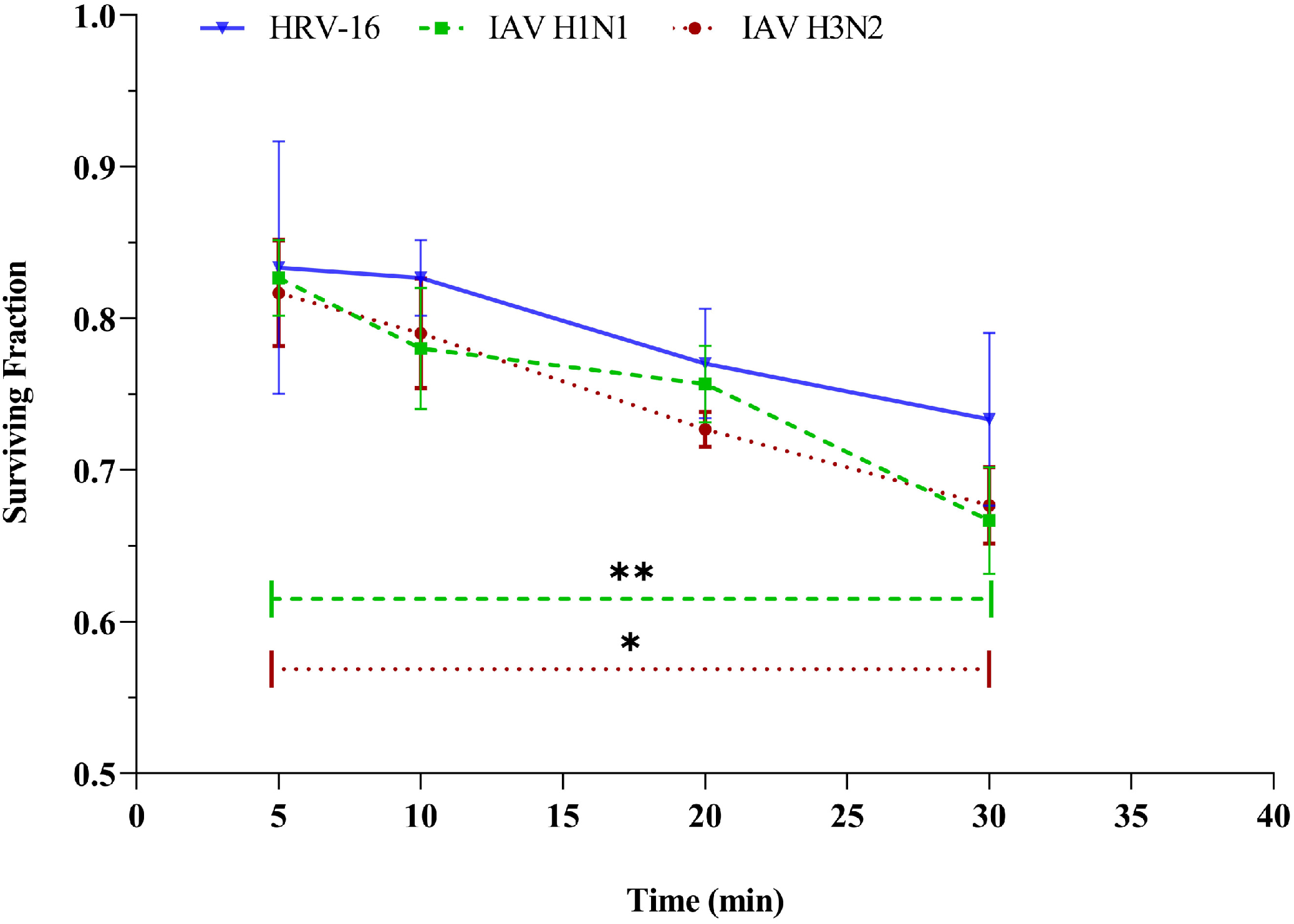
IAV H1N1, IAV H3N2 and HRV-16 SF declines following 30 min of 1-jet Collison nebulization. The 5 min SFs are taken from figure 3. The solid dash and dot lines present the statistical differences of SFs for H1N1 and H3N2 between 5 and 30 min.

## Discussion

The techniques used to generate laboratory bioaerosols simulate the physiological aspects of bioaerosol generation in a controlled environment. These techniques enable greater understanding of the airborne mode in the transmission of respiratory viruses. However, the mechanism of aerosol generation affects the characteristics of the aerosols produced, including their physical and chemical properties, and influences their properties compared to bioaerosols generated from natural sources. We conducted this study to characterize the features of aerosols generated by three commonly used nebulizers, including the 1-jet Collison nebulizer, VMN and HSA. We also tested nebulizer impacts on the stability and infectivity of IAVs H1N1, H3N2 and HRV-16 viruses immediately post-nebulization. Our detailed characterization found that the Collison produced aerosols with CMD of 0.25 µm when derived from a solution of 9 g L^-1^ NaCl concentration. This salt concentration is consistent with the salt concentration of standard cell culture media as the potential suspension for viral particles. The size diameters of the HRV-16 (23) and IAVs (24) are 30 and 80-120 nm, respectively, therefore, any aerosols smaller than this would not be able to carry viral particles, although could carry non-infectious viral fragments. Previous studies reported that a Collison nebulizer produces high concentrations of aerosols which were monodisperse with a MMD between 1-2 µm (25). Based on the particles size distributions generated from a Collison nebulizer, May et al. stated that only 1 percent of the mass of producing aerosols is larger than 5 µm (26, 27). A study conducted by Ibrahim et al. reported that the CMD of the aerosols produced by the Collison nebulizer is between 33 and 38 nm, which is not large enough to carry an Influenza virus particle (> 80 nm) (28). However, fluid physiochemical properties such as viscosity, surface tension (29) and the type and concentration of ions (30) can highly influence the nebulizer aerosol performance. Our results indicated that the physical aspects of aerosols, including CMD and VMD depend on the concentration of ions in solution used. VMN produced aerosols in an airborne size range with the CMD of 0.63 µm derived from a solution of 9 g L^-1^ NaCl, which is larger than most viruses, suggesting that VMN is suited to investigations of airborne virus-laden aerosols in terms of aerosol’s physical size. HSA atomized aerosols were larger than those generated by the two other nebulizers, but were still within the airborne range size, with a CMD of 0.76 µm for 9 g L^-1^ NaCl solution. However, HSA nebulization produced a lower concentration of aerosols in the same running time, followed by 1-jet Collison nebulizer. This property could disadvantage studies that require a higher concentration of aerosols or viral doses. In terms of SF reductions of IAV H1N1and H3N2, the decline in viral viability with HSA was minimal, with an average of 0.89±0.05 and 0.94±0.03, respectively, followed by the Collison and VMN. However, VMN produced the highest SFs of HRV-16 (0.86±0.15), which is a non-enveloped virus, and may suggest that non-enveloped viruses survive nebulization better than enveloped viruses, such as IAV. Our findings showed that the SFs of all three viruses were reduced in samples collected from virus suspensions in the glass of the 1-jet Collison nebulizer after 30 minutes running time and that the loss was highest for enveloped IAVs compared to the non-enveloped HRV-16. Consistent with our findings, it was previously reported that the method of the aerosol production in the Collison jar could damage intact pathogens, effecting the pathogen dose required to identify an infection signal (31). Kim et al. reported a loss of 15% in titre of an enveloped coronavirus (80-160 nm) during Collison nebulization over a period of 30 minutes (32). Conversely, Hermann et al. reported no loss in the viabilities of enveloped porcine reproductive and respiratory syndrome virus (40-80 nm) through 55 min of nebulization using a 24-jet Collison (33). The inconsistency between reported results of viable fractions post-aerosolization is likely due to large variations in methods used for measuring SFs and differences in the size of viruses tested. Experimental protocols which significantly impact results include aerosol sampling and measuring, and the degree of control over temperature and RH. In fact, the Collison nebulizer inherently needs a high volume of virus suspension (10 mL) compared to other nebulizers, which can dilute the titre of initial virus stocks and reduce the aerosolized doses provided. The Collison also generates foam from suspensions with high organic content which could be a barrier for proper aerosolization of virus laden droplets (**Supplementary Figure 1**). Finally, we summarized the advantages and disadvantages of investigated nebulizers in **Table 2**.

**Table 2.**
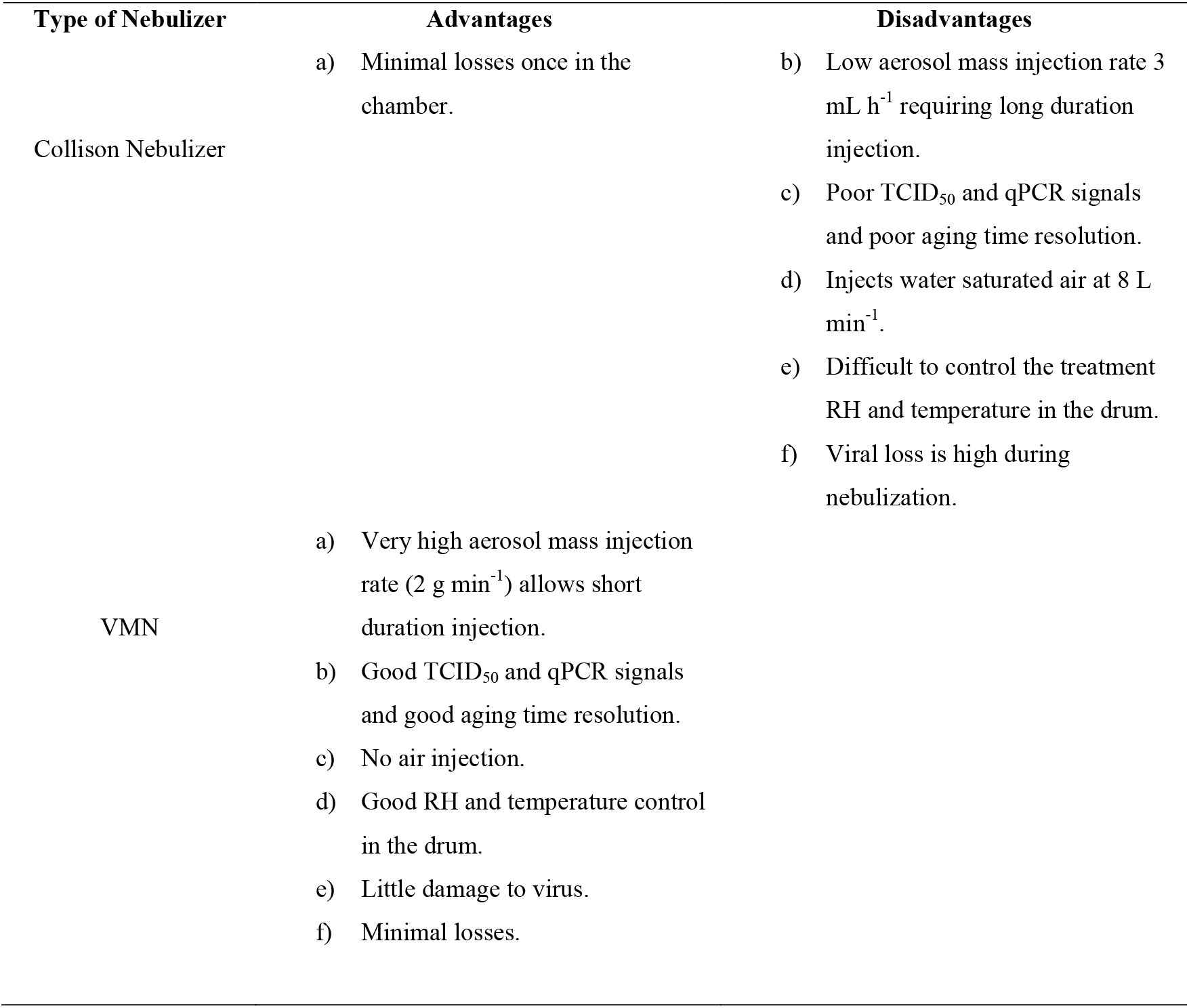
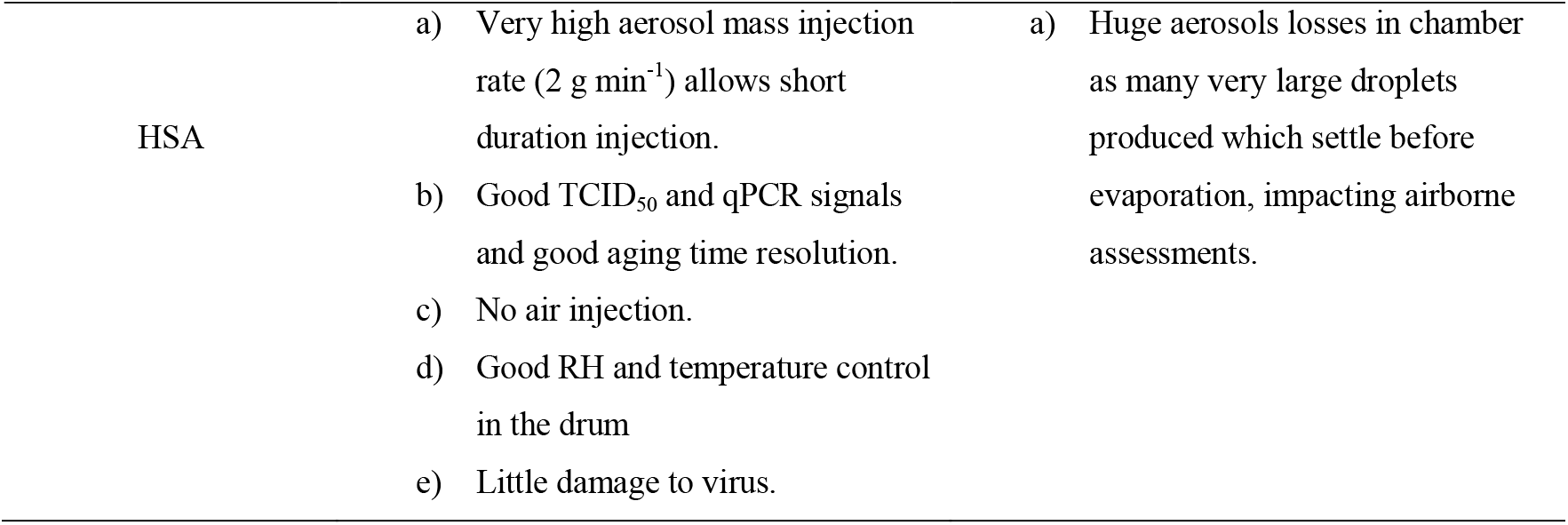
advantages and disadvantages of nebulizers for studies in aerobiology field.

This study was conducted to investigate the characteristics of aerosols generated by commonly used nebulizers using SMPS instrumentation. CMD and VMD of aerosols generated by the Collison placed at 0.25 µm and 2.91 µm, while VMN produced aerosols with CMD and VMD of 0.63 µm and 3.2 µm. HSA generated larger aerosols with CMD and VMD of 0.76 µm and 2.43 µm. VMN generated higher concentrations of aerosols over the same running time compared to the other two nebulizers. SFs of IAV H1N1, IAV H3N2 and HRV-16 decreased post-nebulization, but this was dependent on virus type (enveloped or non-enveloped) and nebulization mechanisms. HSA elicited the least mechanical stress on the investigated IAV strains; however, the gravitational loss of aerosols was higher due to the generation of large size droplets. Our results indicated that VMN is the best nebulizer for infectious disease aerobiology research due to its production of aerosols in a usable size range (200nm-5000nm), higher aerosol mass injection rate (2 g min^-1^) which allows short duration injection and good aging time resolution. The VMN does not inject air, therefore it is suitable for studies that require modulation of RH and temperature control within enclosed chambers.

## Materials and Methods

### Aerosol Size Distribution Measurement

The droplet size distribution produced by each of the nebulizers was back calculated after measuring the dry salt aerosol size distribution produced from a saline solution of known concentration. Aerosols (without addition of viruses) were generated by briefly nebulizing a suspension of 0.05 g L^-1^ NaCl into a 400 L rotating stainless steel drum developed based on the TARDIS-Rotator (34). The drum was first flushed with HEPA-filtered air and the relative humidity (RH) was adjusted to 30%, well below the efflorescence RH of NaCl and the nebulization time limited to ensure generated aerosols dried rapidly and were sampled by the aerosol measurement instrumentation in the dry state. The dry aerosol size distribution (ASD) of aerosols produced by the 1-jet Collison nebulizer, VMN, and HSA were measured using a scanning mobility particle sizer (SMPS 3034, TSI Inc., Shoreview, MN, USA). A dilute sample solution (0.05 g L^-1^) was used for measurements to ensure that the dry aerosols were within the size range of the SMPS which is between 9 to 1000 nm, depending on operating conditions. We then calculated the initial droplet sizes. Subsequently we were able to calculate the dry sizes of aerosols expected from a solution with salt concentration of 9 g L^-1^, similar to a physiological saline solution, using ***Equation 1*** and this size distribution was validated through direct measurement of the portion of that dry size distribution accessible to the SMPS.

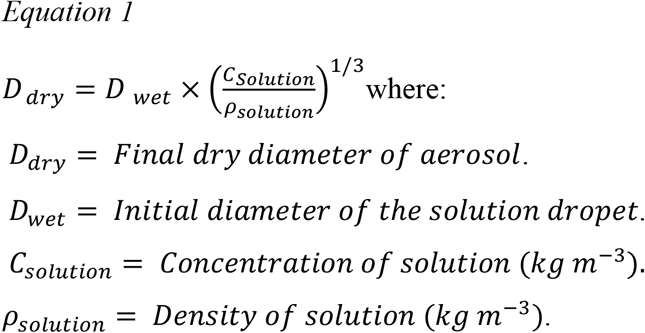

### Virus Propagation

HRV-16 was grown in Ohio HeLa cells in VP-SFM medium (Life Technologies, USA) and propagated according to standard protocols (35) (**Supplementary Text I**). IAV H1N1 and H3N2 were propagated in the allantoic fluid of 10-day old embryonated chicken eggs (36) (**Supplementary Text II**).

### Virus Suspensions

IAV H1N1, IAV H3N2 and HRV-16 virus batches were added to a suspension composed of phosphate buffer saline (PBS) and fetal bovine serum (FBS). The salt and total protein contents were 10 and 8 g L^-1^, respectively, which is approximately comparable to the ratio of salts and proteins found in human respiratory fluid (37). This solution provided an environment with neutral pH that allowed the viruses to remain viable, while attempting to physiologically model the actual composition of human respiratory fluid.

### Experimental Setup for Measuring Viruses Viabilities

**Figure 4**. illustrates the experimental set up for testing the mechanical stress of Collison nebulizer (upper panel), VMN and HSA (lower panel) on the viability of viruses. HEPA-filtered compressed air with a flow rate of 12 L min^-1^ and pressure of 26 to 30 lb in^-2^ was blown into the glass of the 1-jet Collison nebulizer to produce carrier aerosols. Virus-laden aerosols were collected into an SKC Biosampler directly after nebulization in 5 mL PBS. The photographs of experimental set ups also are shown in **Supplementary Figure 2**. The SKC Biosampler (#225-9593, SKC) is an advanced impinger-type air sampler that collects airborne aerosols using a whirling flow of liquid (PBS) and pump flow rate of 12.5 L min^-1^. The SKC swirling flow action is created by drawing air through three 0.630 mm tangential sonic nozzles (38). A HEPA filter was connected to provide required excess air for sampling. Our VMN (**Supplementary Figure 3**) was assembled inside a biosafety cabinet. It consisted of the nebulizer electroform plate, 3 mL reservoir and electrical wires, which were connected to a high frequency circuit inside of the biosafety cabinet. The virus suspension was injected into the nebulizer reservoir via a Luer lock syringe. As the final step, the virus suspension was aerosolized directly into an SKC cup and simultaneously collected into 5 mL PBS by running the SKC pump. The same experimental set up was applied for testing HSA and the virus suspension was nebulized using a 10 mL Luer lock syringe.

**Figure 4.**
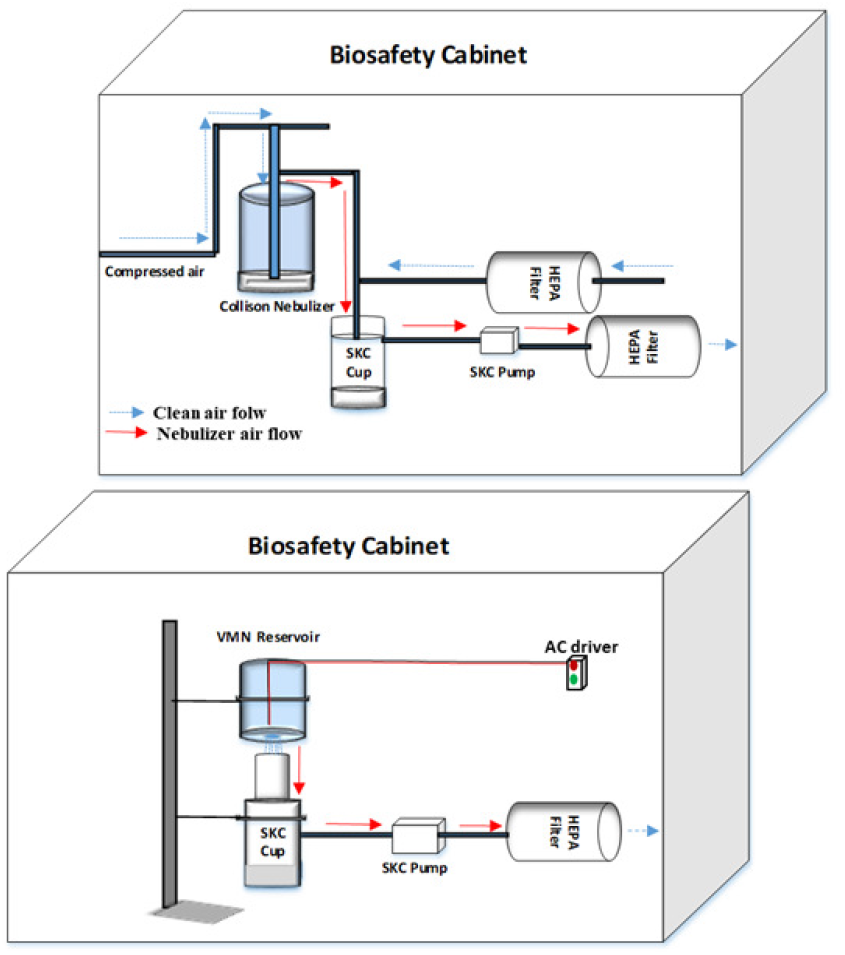
Experimental set up for measuring the effects of nebulization on the viability of respiratory viruses (upper panel: Collison nebulizer and lower panel: VMN and HSA).

### TCID_50_ to Determine HRV-16 Titer

The change in titer of HRV-16 before and after nebulization was quantified using a standard TCID at which 50% of HeLa cells cultured in monolayer demonstrated a cytopathic effect (CPE; TCID_50_) (39). Cells were exposed to replicate 10-fold serial dilutions of the fluid collected for HRV-16 in the BioSampler for four days at 34 °C. CPE was then identified for each well in the TCID_50_ assay using 0.01 % crystal violet solution mixed with 60% ethanol and 40% methanol.

### Plaque Assay to Determine IAV H1N1 and IAV H3N2 Titers

IAV H1N1 and H3N2 infectivity titers were measured by plaque formation on confluent monolayers of Madin-Darby canine kidney (MDCK) cells cultured in RF10 medium (40). Once cells in 6 well plates were confluent, media was replaced with 135 µL RPMI containing 10-fold serial dilutions of recovered sample. Plates were incubated (37 °C, 5% CO_2_) for 45 min, and each 15 min gently rocked to distribute the virus evenly and prevent drying of the cell monolayer. Cells were then overlayed with 3 mL per well of L15 media mixed with 0.1% trypsin and 1.8% (w/v) agarose. Once set, plates were incubated (37 °C, 5% CO_2_) for 3 days. Plaque formation was visualized by staining the monolayer with 0.01% crystal violet solution, and number of plaques were counted as a measure of virus infectivity. Experimentation was conducted according to health risk assessment, QUT biosafety committee approval (Approval number:1800000969).

### Surviving Fraction and Statistical Analysis

Surviving Fractions (SFs) of IAVs (H1N1/H3N2) and HRV-16 were calculated using ***Equation 2***. RNA was used as a natural tracer of dilution based on the expectation that RNA was present in the same ratio to total virus in the nebulizer before aerosolization and after sample collection (39). Viral RNA was extracted from nebulizers and SKC BioSampler samples by QIAamp Viral RNA Mini Kit (Qiagen, USA) and its concentration quantitated using Qubit High Sensitivity RNA kit (Thermo Fisher Scientific, USA).

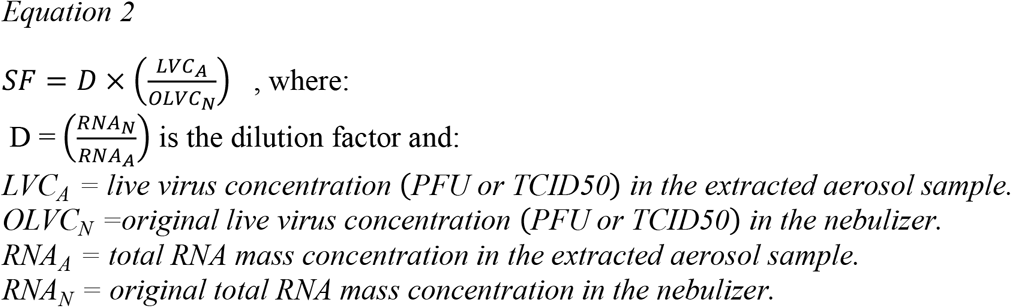

The concentrations of extracted RNA (ng mL^-1^), PFU mL^-1^ (IAV H1N1 and IAV H3N2) and TCID_50_ mL^-1^ (HRV-16) for each experiment operated by 1-jet Collison, VMN and HSA are presented in (**Supplementary Table 1). Supplementary Table 2** also shows the concentrations of extracted RNA (ng mL^-1^), PFU mL^-1^ (IAV H1N1 and H3N2) and TCID_50_ mL^-1^ (HRV-16) for each experiment operated by 1-jet Collison after 10, 20 and 30 minutes nebulization. All experiments were conducted in biological triplicate. A One-way analysis of variance (ANOVA) was performed to compare SFs of viruses between nebulizers. P value less than 0.05 was considered statistically significant.

## Supplementary Data

Supplemental material for this article may be found at…

## Acknowledgements

The research described here was sponsored by Australian Research Council (grant number DP170102733). We thank Dr. Kirsty R short for allowing us to propagate influenza A viruses in her laboratory.

